# Scalable method for micro-CT analysis enables large scale quantitative characterization of brain lesions and implants

**DOI:** 10.1101/2020.03.12.989871

**Authors:** David B. Kastner, Viktor Kharazia, Rhino Nevers, Daniela A. Astudillo-Maya, Greer M. Williams, Zhounan Yang, Clay Smyth, Luca Della Santina, Dilworth Y. Parkinson, Loren M. Frank

## Abstract

Anatomic evaluation is an important aspect of many studies in neuroscience; however, it often lacks information about the three-dimensional structure of the brain. Micro-CT imaging provides an excellent, nondestructive, method for the evaluation of brain structure, but current applications to neurophysiological or lesions studies require removal of the skull and hazardous chemicals, dehydration or embedding, limiting their scalability and utility. Here we present a protocol using eosin in combination with bone decalcification to enhance contrast in the tissue and then employ monochromatic and propagation phase-contrast micro-CT imaging to enable the imaging of brain structure with the preservation of the surrounding skull. Instead of relying on descriptive, time-consuming, or subjective methods, we develop simple quantitative analyses to map the locations of recording electrodes and to characterize the presence and extent of brain lesions.

## Introduction

Evaluating brain structure is a critical confirmatory step for much of systems neuroscience. This often occurs through histological sectioning and light-microscopic analysis. Although such a technique provides a high-resolution view of brain structures, it is time-consuming, difficult to scale for high-throughput analyses, and, perhaps most problematic, it destroys the three-dimensional structure of the brain. Additionally, histological processing is often accompanied by descriptive analyses and not quantitative evaluation. Newer tissue clearing methods^1,2^ maintain the three-dimensional structure of the brain for high resolution imaging, but remain expensive, labor intensive, and not well suited to evaluate lesions or electrode localization, mainstays of systems neuroscientific inquiry. Micro-CT presents an excellent middle ground for evaluating large-scale three-dimensional brain anatomy while maintaining the original structure^3-9^, providing an excellent substrate for subsequent quantitative analyses^10-15^.

The majority of studies that visualize neuroanatomy using micro-CT have not evaluated the technique for quantifying brain lesions and electrode placement, instead focusing on tumors^4,5,16^, cavernous malformations^10,17^, disease pathology^15,18-20^ and cellular and subcellular visualization^11-13,21,22^. A recent protocol was developed for micro-CT imaging of lesions and electrode placement^8^ providing an elegant proof of principle of the use of micro-CT for systems neuroscience. However, that protocol, although clear and well described, contains many involved steps, some of which require expensive and hazardous elements, including osmium. It also removed the skull prior to imaging, leaving out valuable information and removing a structural support for neural hardware. Furthermore, in using osmium, it becomes far more challenging to process the brain for subsequent standard histologic techniques, requiring a decision to be made between either imaging the full three-dimensional volume, or using standard histologic processing.

Phase contrast micro-CT has proven capable of solving all of the above-mentioned problems. Utilizing phase contrast techniques to process micro-CT images generates sufficient contrast in the brain^3,6,7,9,15,23^ without, heavy metals or complex protocols. In addition, brain structures can be resolved with the overlying skull intact^9^, histology can be performed on the same samples following the CT^5,11,14-16,20,21,24^, and laborious embedding of the tissue is not necessary^5^. However, phase contrast processing, to the best of our knowledge, has not been applied to brains with recording electrodes in place, nor has it been used to evaluate brains lesioned for behavioral studies. Moreover, quantitative approaches to electrode localization or lesion quantification using phase contrast processing have not been presented.

Here we present a scalable and inexpensive protocol utilizing monochromatic X-rays and propagation phase contrast to image and quantify brain lesions and electrode locations within the brain. We adapt decalcification techniques^25,26^ to image the brain with critical parts of the skull intact. Furthermore, we avoid any dehydration or embedding of the tissue, and enhance contrast to the post-mortem brain using eosin as an additional contrast agent^27,28^. The resulting protocol allows for evaluation of brain structure relative to the coordinates determined by skull sutures and permits further processing with standard histologic techniques.

## Material and Methods

### Animals

All experiments were conducted in accordance with University of California, San Francisco Institutional Animal Care and Use Committee and US National Institutes of Health guidelines. Datasets were collected from male Long-Evans rats that were fed standard rat chow (LabDiet 5001). Long-Evans rats used for the lesioning were either of a wild-type genotype or of a *Fmr1* KO genotype^29^ and were bred at Medical College of Wisconsin, the remaining rats were purchased from Charles River. We did not observe any gross anatomical differences between the genotypes; therefore, we combined all brains for the analysis of the lesions.

### Surgery

Anesthesia was induced using ketamine, xylazine, atropine, and isoflurane. For the animals that underwent surgery, they were either implanted with a multielectrode microdrive, or underwent lesioning of the dorsal and intermediate hippocampus. For the microdrive implant, craniotomies and accompanying durotomies were placed at +2.6 ML and −4.0 AP for the dorsal hippocampal canula and +4.7 ML and −5.7 AP for the intermediate hippocampal canula. Values are relative to bregma. Screws were imbedded into the skull to support the acrylic casing that secured the microdrive onto the skull of the rat. For the lesion rats, craniotomies were drilled bilaterally to allow for access to all 24 injection sites (Table 1) and the injection needle (33 gauge) was inserted through the dura. For each lesion site, the needle was lowered 0.1 mm below the target coordinates and then raised back to the target prior to injection. 120 nl of NMDA (10 mg/ml) was injected at each site at a rate of 0.1 μl/min. The needle was left at the site for 2 minutes after finishing the injection. For the rats that underwent control surgeries, everything was the same as the lesion rats, except no NMDA was injected, and the needle did not dwell at the hypothetical injection site. The animals that underwent the lesion surgery (either with or without injection of the NMDA) received Diazepam 10 mg/kg intraperitoneally post-operatively.

**Table 1:** Stereotaxic coordinates of NMDA infusions to produce lesions within the hippocampal formation. Adapted from Kim and Frank^47^. The coordinates are given for a Long Evans rat skull which is leveled so that bregma and lambda line in the same horizontal plane. AP: anteroposterior, ML: mediolateral, DV: dorsoventral. The AP and ML coordinates are measured from bregma. DV coordinates are measured from the dural surface above the brain.

| AP (mm) | ML (mm) | DV (mm) |
| --- | --- | --- |
| -2.8 | ±1.4 | -3.0 |
| -3.3 | ±2.4 | -3.0 |
| -4.1 | ±1.8 | -2.8 |
| -4.1 | ±3.4 | -2.8 |
| -4.8 | ±2.0 | -2.8 |
| -4.8 | ±4.2 | -3.1 |
| -5.5 | ±2.6 | -3.0 |
| -5.5 | ±3.6 | -2.9 |
| -5.5 | ±5.0 | -3.5 |
| -5.5 | ±5.0 | -5.5 |
| -6.2 | ±4.0 | -3.4 |
| -6.2 | ±5.4 | -4.4 |

### Electrophysiological recording

Tetrodes were individually moved toward the hippocampus daily. Physiological data, digitized at 30 kHz, and was recorded using the SpikeGadgets data acquisition system and the Trodes software package (SpikeGadgets).

### Processing for micro-CT

Prior to perfusion, animals were anesthetized with isoflurane, and then euthanized with pentobarbital. Animals were perfused intracardially with 100 mL of phosphate buffer solution (PBS), followed by 250 mL of chilled 4% paraformaldehyde in PBS. The skull was dissected out with the brain intact, with most of the facial bones trimmed off, and all soft tissues removed. A 5-mm-wide opening was made using a rongeur on the ventral side along the midline (optional), for improved diffusion of fixative. The sample was then placed in 50mL of the same fixative for 48 hours at 4°C.

Subsequently, the sample was rinsed in PBS with 0.05% sodium azide (PBS-azide) and decalcified by placing it in a container with 250 mL of 8% EDTA (Sigma-Aldrich) for 6-7 days at room temperature on an orbital shaker. The solution was replaced once midway through the decalcification.

After rinsing again in PBS-azide, the decalcified skull was further trimmed to a final size on both anterior and posterior sides using a razor blade. Care was taken to avoid compressing or otherwise distorting the brain. The trimmed sample was placed in 50mL of 5% Eosin Y (Sigma-Aldrich) for 6-7 days at 4°C on an orbital shaker. The solution was replaced once after 48 hours. The sample was then rinsed in PBS and stored in 50mL PBS-azide at 4°C prior to micro-CT scanning.

For Nissl staining, the brain (with skull removed) was cut using a Vibratome 3000 (Leica Biosystems); 50-80 um thick coronal sections were collected into a 24-well dish with PBS-azide and stored at 4°C. Sections were mounted on adhesive slides, air-dried, and stained using a traditional cresyl violet staining protocol.

We confirmed the benefits of the eosin staining by processing and imaging a brain without eosin and found that the eosin provided substantial improvements in contrast (data not shown).

### Micro-CT scanning and image reconstruction

Scanning was done at beamline 8.3.2 (hard X-ray micro-tomography) at the Advanced Light Source (ALS) at Lawrence Berkeley National Laboratory (Fig S1). All scans were carried out at room temperature.

For the brain with the implanted microdrive, the brain anterior and posterior to the hippocampus were removed to decrease the width of the brain to minimize the field of view for the imaging. Furthermore, as many as possible of the skull screws were removed to minimize interference. As the microdrive was still attached to the skull via acrylic and remaining skull screws, it provided the support for the brain. Consistent with a previous protocol^5^ the brain was out of solution for the scanning. X-rays passed through the sample along the A-P/M-L axes as the sample rotated.

For the rest of the brains, the sample was placed snuggly in a conical tube and was not in solution (Fig S1), consistent with a prior protocol^5^. X-rays passed through the sample along the M-L/D-V axes as the sample rotated. This is the narrowest dimension of the brain, thereby allowing for the minimal field of view and finest pixel size (see Table 2), as limited by the number of pixels on the available detector at the beamline that can accommodate the full field of view of the brain.

**Table 2:** Scanning parameters and reconstruction parameters for Tomopy for all of the brains. The lesion brains include all of the brains described in Fig 3, including both the brains that underwent control and lesion surgery. The implanted brain was processed in two ways, one two resolve the brain anatomy and the other to maximally resolve the electrodes. The x-ray beam is only so wide; therefore, to scan an entire volume multiple tiles in the vertical dimension were taken and then the tiles were manually stitched together. The lesion brains were imaged from the striatum to the cerebellum to focus on the brain regions of interest and to save time.

|  | full brain (Fig 1&2) | lesion brains (Fig 3) | implant brain: electrodes (Fig 4) | implant brain: anatomy (Fig 4) |
| --- | --- | --- | --- | --- |
| energy | 25 keV | 25 keV | 25 keV | 25 keV |
| angles | 1313 | 1313 | 1313 | 1313 |
| exposure | 125 ms | 100 ms | 125 ms | 125 ms |
| pixel size | 7.205 $\mu\text{m}/\text{pixel}$ | 7.84 $\mu\text{m}/\text{pixel}$ | 7.72 $\mu\text{m}/\text{pixel}$ | 7.72 $\mu\text{m}/\text{pixel}$ |
| scintillator distance | 540 mm | 540 mm | 540 mm | 540 mm |
| # tiles | 32 | 8 | 4 | 4 |
| ~scan time | 150 min | 32 min | 25 min | 25 min |
| outlier removal in 1d | true | true | False | true |
| ring removal | polar ring | polar ring | Fourier-wavelet | polar ring |
| max width for ring removal | 100 | 100 | 100 | 200 |
| min value of image to filter | 0.5 | 0.6 | -3000 | 0.6 |
| max portion of image to filter | 1.2 | 1.2 | 3000 | 1.2 |
| manual centering | false | false | true | true |
| phase retrieval | true | true | false | true |
| alpha regularization | 0.0005 | 0.0005 | — | 0.0005 |

Monochromatic X-rays were generated using a multilayer mirror monochromator. Energy of 25 keV was used. The flux is approximately ∼10^5^ hν/sec/μm^2^. The source size is ∼220 x 30 μm full width at half maximum, and the maximum beam size at the sample is ∼40mm x 4.6 mm. The sample was placed 54 cm from the scintillator (Fig S1) to enable the maximal usage of propagation phase contrast imaging available at the beamline (because of its small hutch, this is the maximal sample to scintillator distance for this beamline). For each scan, a total of 1313 projection angles with an exposure time of 100 – 125 ms each were used (Table 2). Detection was accomplished with a 0.5 mm thick YAG:Ce scintillator, a Nikon zoom lens, and a PCO.Edge sCMOS detector. The zoom was set to resolve 7 μm lines and spaces (14 μm period) on a test pattern in 2 dimensions. We did not measure precisely the 3-dimensional resolution as it was not necessary for the conclusions of this study, but our results are consistent with an approximately 14 μm isotropic resolution. Radiation dose was not calculated for this study as we did not detect any detrimental effects of the scanning at the resolution used (data not shown).

The images were reconstructed using tomopy using a script designed at the ALS (https://bitbucket.org/berkeleylab/als-microct-python/src/master/tomopy832/tomopy832.py). See Table 2 for the parameters used for the reconstruction of the various brains. Multiple tiles were manually stitched together to create the final imaged volume (Table 2). It is the stitching together of the multiple tiles that causes the striped patterning see in (Fig 1 & Vid 1). Phase retrieval was accomplished using the standard Paganin method^30^, as implemented in tomopy.

**Figure 1:**
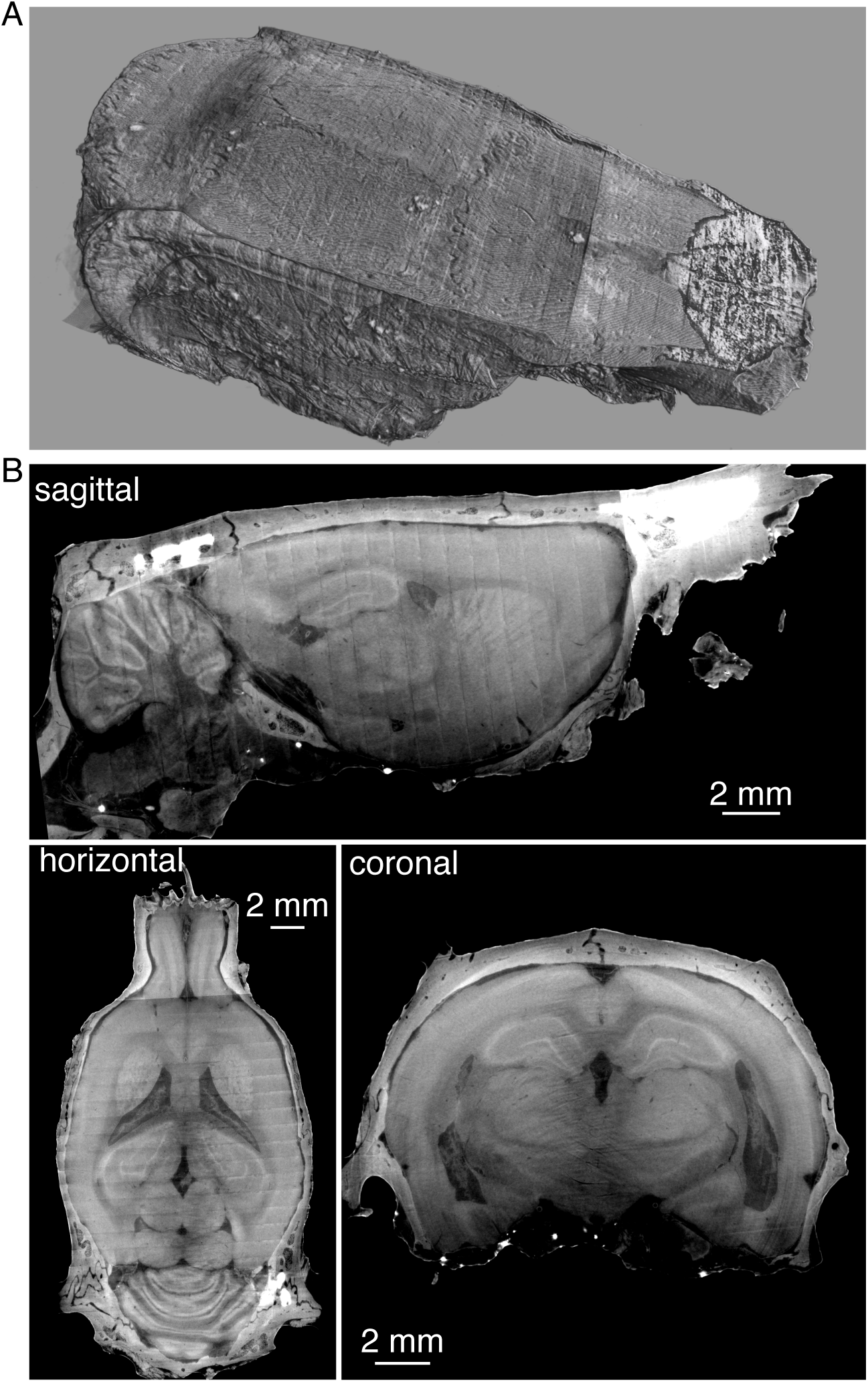
micro-CT image of the intact brain with overlying skull. A) Three-dimensional representation of the overlying skull. B) The same brain and skull from A, virtually sliced along the sagittal plane (top), horizontal plane (bottom right) and coronal plane (bottom left).

### Image visualization and analysis

Fiji^31-33^ was used to visualize virtual slices of the brain and to create a mask of the hippocampal region for the lesion analysis using the Segmentation Editor tools. Imaris (Oxford Instruments) was used for three-dimensional reconstruction of the brain. Slicer-3D 4.10.2 (https://www.slicer.org/)^34,35^ was used to register and align the brains (http://hdl.handle.net/1926/1291) by finding the linear transformation (rotation and translation) to place the entirety of the imaged brains into a shared coordinate system. The alignment of the brains was not restricted to align any specific region of the brain (such as just the hippocampus) and rather worked to best align the whole brain and skull.

For the lesion analysis we were interested in understanding the similarity of the hippocampal formation between brains as that was the target of the lesioning. Therefore, we created a mask for the left and right hippocampal region. The mask was intentionally generous to ensure that the entire hippocampal formation was included for all brains. For each virtual slice we calculated the similarity between all pairs of brains using the structural similarity metric (SSIM)^36,37^. For SSIM we used a gaussian smoothing filter for the images with a sigma of 50 pixels. This allowed for better comparison between the brains due to inherent differences in the exact positioning of the brain structures. A dendrogram that grouped lesioned and control brains was calculated using single-linkage hierarchical clustering over the average SSIM for all pairs of brains. The similarity matrix was subtracted from 1 to create a dissimilarity matrix as the input to the hierarchical clustering.

### Theta phase offset calculation

We also calculated the phase of the endogenous ∼8 Hz theta rhythm at each electrode so that differences in phase across electrodes could be related to electrode locations. The local field potential (LFP) was defined as the voltage from each tetrode after being low-pass filtered below 400 Hz and down-sampled to 1.5 kHz. To extract theta oscillations, the LFP data were band-pass filtered with a Parks-McClellan optimal equiripple FIR filter (pass band between 4 and 10 Hz, stop bands below 3 Hz and above 11 Hz) applied in both the forward and reverse directions to prevent phase distortion^38^. Instantaneous theta phase was calculated as the angle of the Hilbert-transformed theta oscillation.

Analysis of theta phase was limited to time periods when the rat was running, which was defined as a body speed greater than 10 cm/s^38^. Instantaneous phase differences between pairs of tetrodes were measured as the angle between the complex phase vectors for each tetrode. For each running period, the average pairwise phase difference was calculated as the circular mean over time of the angle between phase vectors, and the average phase offset over the entire behavior session was calculated as the circular mean of individual average phase differences for each running period.

## Results

### Brain structure with intact skull

We sought to develop a protocol that does not require osmication, dehydration of the tissue, nor embedding in resin or paraffin, but could still provide sufficient contrast to image brain structures in the presence of electrodes or lesions. We wanted to avoid osmication and other heavy metals as they can add laborious steps to the processing of the brain and can often add significant expense. We wanted to avoid dehydration of the tissue and embedding in resin or paraffin as these techniques often lead to substantial shrinkage of tissue^39,40^. We therefore adapted protocols using eosin^27,28^ to provide enhanced contrast within the brain, and imaged the brain using monochromatic X-rays at the Advanced Light Source at Lawrence Berkeley National Lab.

Typically, the skull is removed to image the brain, as it provides a barrier for X-ray transmission. However, the skull provides the reference frame for the stereotactic coordinates utilized to target specific brain structures while the animal is alive. Removing the dorsal skull therefore discards a piece of information critical for evaluating and refining anatomical targeting. Furthermore, the skull also provides a rigid structure to prevent alterations to the brain dimensions due to histological processing. Therefore, in addition to processing the brain with eosin, we adapted decalcification techniques to leave most of the skull in place but allow for more transmitted X-rays to better resolve brain anatomy. Leaving the skull intact also allowed for a direct confirmation that the brain maintained its integrity through the processing and imaging (see Anderson & Maga^41^ for a counterexample showing brain shrinkage relative to the skull).

With our method we are able to image the whole rat brain with the overlying skull (Fig 1 & Vid 1). The eosin combined with phase contrast processing provides sufficient contrast to the brain to allow for the visualization of many different brain regions. Given that the imaging provides a three-dimensional volume, we can virtually section the brain to any given region of interest along any plane of interest (Fig 1B). Having the skull intact allows for setting the frame of reference of the brain as one would in surgery by leveling bregma and lambda.

An advantage of using eosin as a contrast agent is that eosin is commonly used stain for histology, and therefore still allows for classic histologic imaging following the micro-CT. Therefore, using the same brain, following the micro-CT, we sectioned the brain and stained it with Cresyl Violet to demonstrate the Nissl substance in the neurons. Nissl-stained brain sections exhibited well preserved morphology and can be used as anatomic reference to corresponding micro-CT virtual slices (Fig 2). The Nissl-stained brains also confirmed the preservation of relevant brain structure through the preparation and imaging steps.

**Figure 2:**
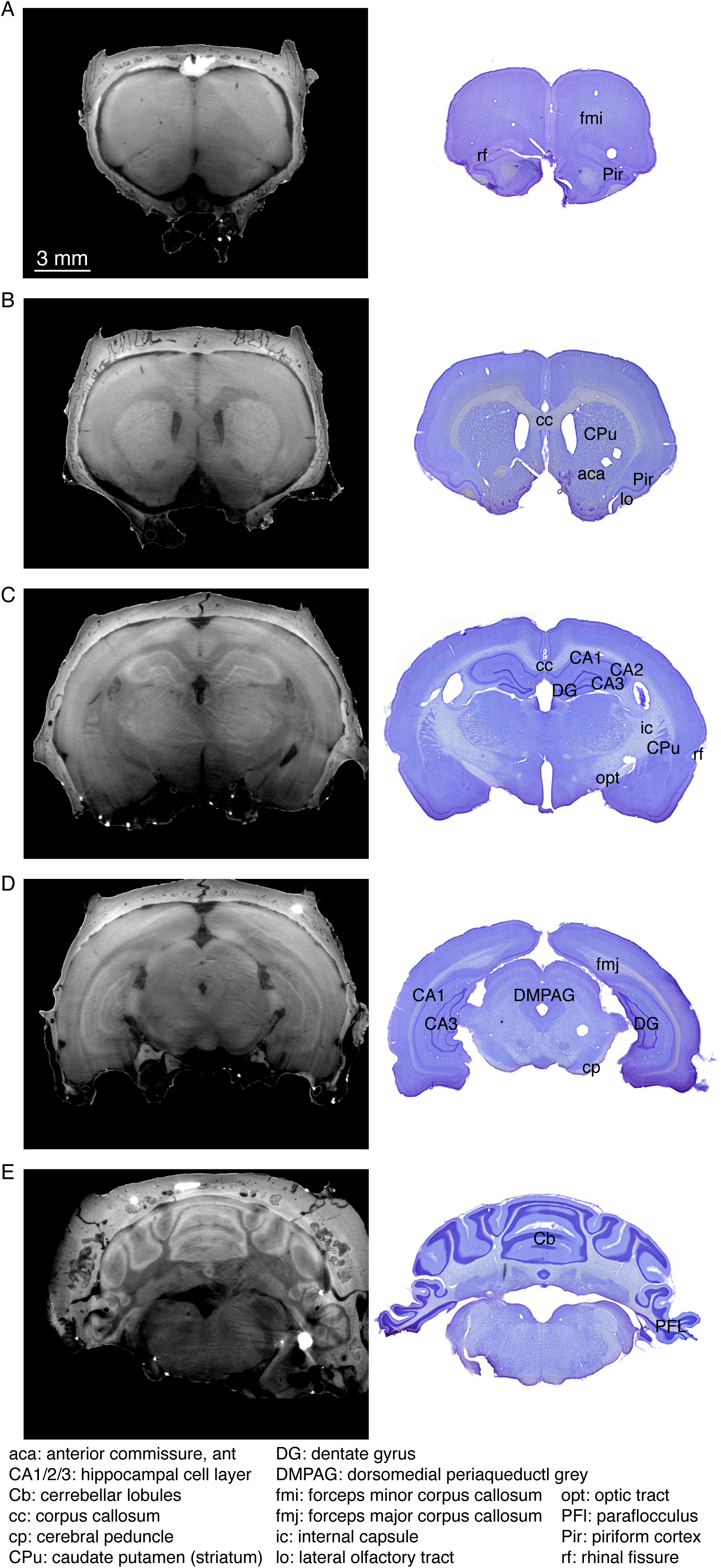
Comparison of virtual slices to standard histological sections following Nissl (Cresyl Violet) stain. The same brain was utilized for both imaging modalities. A-E show different coronal slices along the anterior-posterior axis. Within each slice, sample brain regions and anatomic landmarks that can be visualized both in the CT and Nissl stain are indicated using the brain atlas by Paxinos and Watson^46^. Scale bar in A reflects scaling for all panels.

### Lesion quantification

Lesions provide a useful perturbation for evaluating the necessity of brain regions for behavior. However, validating a lesion using sectioning, staining and imaging presents a challenge for quantification, often leading to more descriptive analyses. Therefore, we utilized our protocol to quantitatively evaluate the presence and extent of hippocampal lesions.

The CT protocol enables clear visualization of the lesioned area of the brain (Fig 3A). Quantification of the lesion could be done by image segmentation, as was done with previous micro-CT processing^8^; however, image segmentation is laborious and prone to subjectivity. Therefore, we sought a more automatable and objective approach for quantifying the presence and extent of lesions.

**Figure 3:**
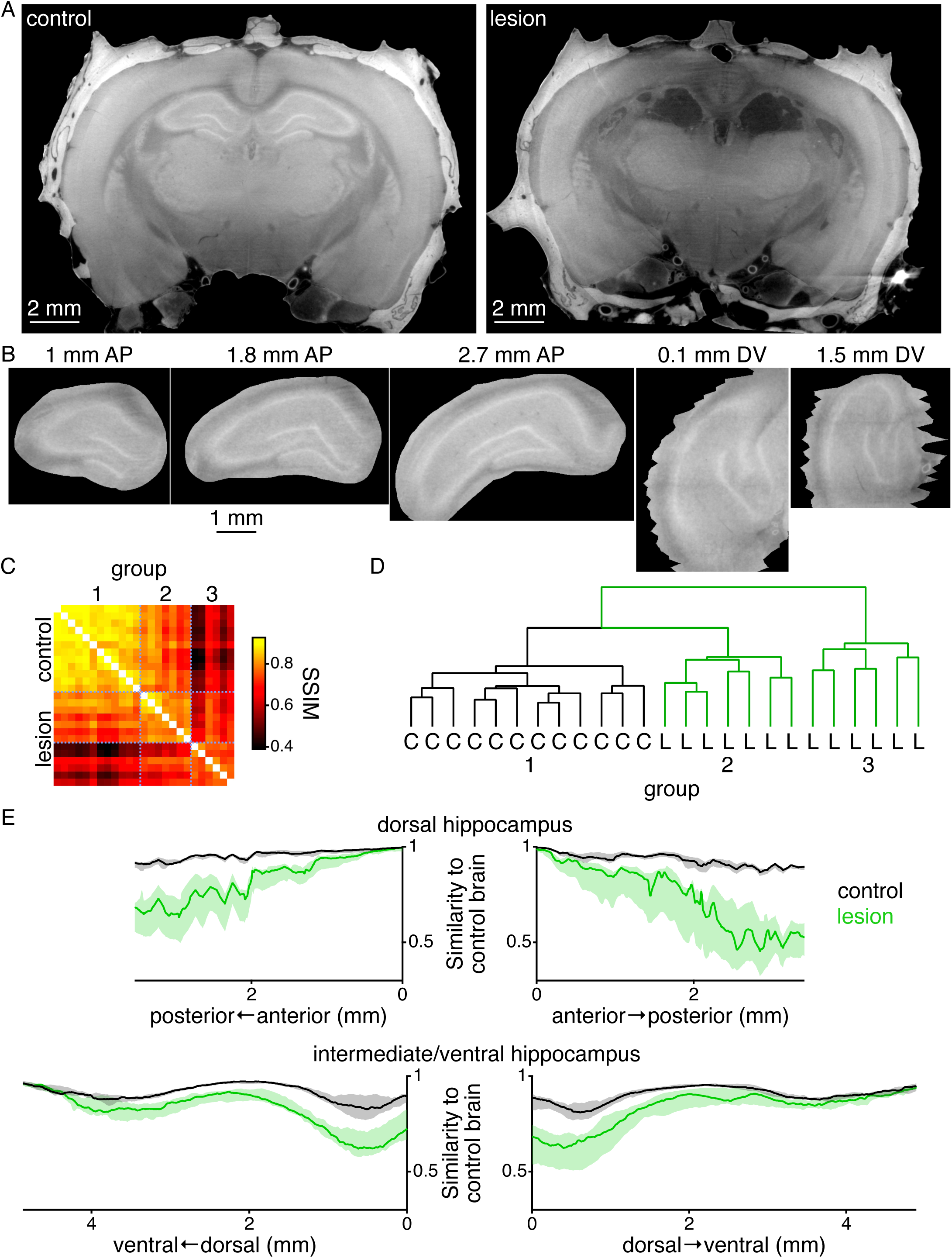
Visualization and quantification of hippocampal lesions. A) Virtual coronal slice through example brains that underwent control surgery (left) and hippocampal lesion surgery (right). B) Example virtual slices of the hippocampus following aligning all brains and masking the hippocampus. Dorsal hippocampus is shown along the anterior-posterior axis and intermediate hippocampus is shown along the dorsal-ventral axis. Jagged edges seen in the dorsal-ventral slices reflect the fact that the mask was created and interpolated along the anterior-posterior axis. Scale bar applies to all images within this panel. C) Pairwise structural similarity metric (SSIM) between all pairs of brains. Each pixel represents the value of the SSIM averaged across all virtual slices bilaterally throughout the dorsal hippocampal formation and the most dorsal part of the intermediate hippocampal formation. Matrix is ordered based on the hierarchical cluster from panel D as indicated by the group labels at top. Dotted blue lines demarcate the major groups from panel D. Left axis shows which are the control and lesion brains. D) Dendrogram showing the results of the hierarchical clustering of all brain based on the similarity matrix from panel C. Brains labeled as either control (C, black lines) or lesion (L, green lines). E) Median (+-interquartile range) similarity of control (black) and lesion (green) brains to nearest intact brain as a function of anatomical location. Top left and right plots show the similarity metric calculated along anterior-posterior axis of the right and left dorsal hippocampus. Bottom left and far right plots show the similarity metric calculated along the dorsal-ventral axis of the right and left intermediate/ventral hippocampus, respectively.

We processed and collected micro-CT images of 25 rat brains—12 rats underwent control surgery and 13 rats underwent hippocampal lesion surgery (see Material and Methods). We aligned all brains to a common reference and created a mask to extract the three-dimensional region of the hippocampus across all brains (Fig 3B). We calculated the similarity of all pairs of brains for each virtual slice using SSIM (Fig 3C), a similarity metric designed to capture the perceptual similarity between images^36,37^. We clustered the brains using hierarchical clustering based on the average pairwise similarity matrix across the bilateral dorsal hippocampal formation and bilateral dorsal-most portion of the intermediate hippocampal formation (the portion of the hippocampal formation that was lesioned (Table 1)). The lesion and control brains separate into different groups by the clustering (Fig 3D).

To visualize the extent of the lesion, for all brains and virtual slices we found the most similar control brain and then plotted those values for the group of control and lesion brains across the dorsal and intermediate/ventral hippocampus for both the left and right sides of the brain (Fig 3E). Across the entire extent of the dorsal hippocampus, and the initial part of the intermediate hippocampus the two groups of animals were clearly separate. Towards the later part of the intermediate hippocampus and into ventral hippocampus the brains were indistinguishable, both being similar to control brains. This is consistent with the intended location of the lesions, since far intermediate and ventral hippocampus were not targeted (Table 1).

### Electrode localization

Beyond just providing potentially useful landmarks, the skull also provides a stable structure to support a microelectrode drive, thereby enabling minimal disruption of electrodes during processing and scanning. Therefore, we evaluated if we could image functional electrodes implanted in the brain and assess their location relative to brain structures. Independently moveable tetrodes were implanted above dorsal and intermediate hippocampus and recordings were taken as animals explored a spatial environment. Subsequently the animal was sacrificed, and the skull and brain prepared for imaging with the microdrive remaining on the skull to minimally disrupt the electrodes. The brain, with skull and microdrive intact, underwent imaging with micro-CT.

The electrodes were easily visualized, and the resolution was sufficiently fine to distinguish the four individual electrodes that comprised each tetrode wire (Fig S2A). Although the hardware used to stabilize the microdrive on the skull degraded the imaging quality somewhat, there remained sufficient signal to resolve the pyramidal cell layer of the hippocampus (Fig 4A). All that was required to identify individual electrodes was a visual comparison of the layout of the canula of the drive and the arrangement of the electrodes from the micro-CT (Fig 4B).

**Figure 4:**
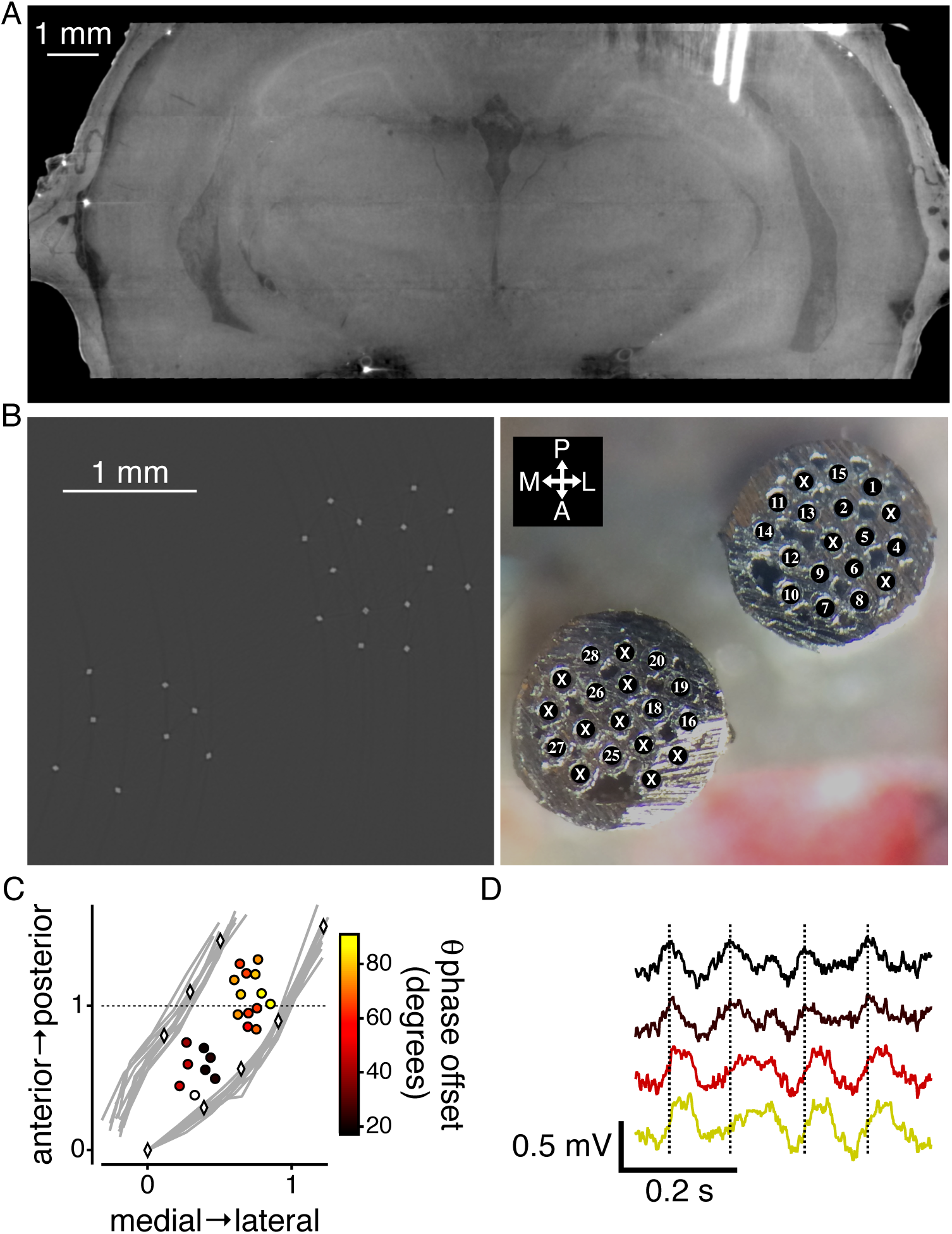
Visualization and quantification of electrode localization. A) Virtual coronal slice through a brain with 22 tetrodes implanted, two of which can be visualized in the slice. B) Virtual horizontal slice through the tetrodes (left). This image was processed to highlight the tetrodes (see Table 2 for parameters). Image of the bottom of the canula of the drive implanted into the brain (right) with the locations of the different tetrodes numbered. Xs indicate locations that did not have tetrodes. C) Two-dimension coordinate system defined by the most anterior part of the hippocampal cell layer (Fig S2B) and the lateral part of hippocampal cell layer at the point of intermediate hippocampal fusion (Fig S2B). Left-most gray lines show the locations of the medial part of dentate gyrus cell layer for 12 different brains. Rightmost gray lines show the locations of the lateral part of the hippocampal cell layer. Diamonds demarcate the same points for the electrode implanted brain. Circles represent the location of the electrode implanted into the brain (one tetrode was utilized as reference and was therefore not included in this plot). Colors within the circles show the mean phase offset of the theta rhythm in the local field potential relative to the electrode denoted by the white circle. D) Example local field potential recordings from electrodes from 4 different tetrodes with traces colored by the phase offset. Black, top, trace is the reference electrode. Vertical dotted lines indicate the peak of the theta rhythm in the reference (top) trace.

As with the lesions, here, with the electrodes, the goal is not just to visualize, but to also extract relevant quantitative information. Again, it would be possible to perform image segmentation for the hippocampal cell layer, but that would be quite laborious and subjective at the most critical parts due to the large signal produced by the electrodes. Instead we sought to extract coordinates that could form the basis of a systematic two-dimensional space on which we could place the locations of the electrodes. For this analysis, we were most interested in a two-dimensional space for the anterior-posterior and medial-lateral axis, since the electrodes were moved over the course of the recording, thereby making the dorsal-ventral coordinates of the CT not representative of the recording location.

We took advantage of the control brains that we scanned for the lesion cohort (n=12) and utilized those brains to define a consistent two-dimensional coordinate system. As the origin of the space we utilize the center of most anterior portion of the hippocampal CA1 cell layer (Fig S2B). And then as point 1,1 we utilized the lateral extent of the hippocampal (CA1/2/3) cell layer at the point where the dorsal and ventral portions of the hippocampus fuse to form intermediate hippocampus (Fig S2B). We chose these points as they were easily visible in all brains and would be likely visible in an electrode implanted brain as well. Within this space, we then plotted, for each brain, sample locations of the lateral aspect of the hippocampal cell layer and the medial aspect of the dentate gyrus cell layer from anterior to posterior (Fig 4C&S2B). Across all brains the locations were very similar.

We then aligned the brain with the implanted electrodes to the same reference brain as the lesion cohort and normalized the space by locating the equivalent points. In this normalized space, the lateral aspect the hippocampal cell layer and the medial aspect of the dentate gyrus cell layer fell within the range of the control brains, confirming the uniformity of the space. This then allows for the placement of the anterior-posterior and medial-lateral location of the electrodes (Fig 4C).

As the goal of localizing the electrodes is to inform the physiology recorded on those electrodes, we took advantage of the finding of theta phase offset in the local field potential as the hippocampus evolves along its septo-temporal axis^38,42^. Within this two-dimensional space we confirmed a phase offset of the theta rhythm as the hippocampus progress from dorsal to intermediate (Fig 4C&D) (see Materials and Methods). Furthermore, this technique provides a straightforward reference to measure and discover further axes along which such things could vary.

## Discussion

We have presented a cost-effective and scalable protocol for utilizing micro-CT to image postmortem brain structure with the overlying skull (Fig 1 & Vid 1). The protocol allows for the quantification of lesions (Fig 3) and localization of electrodes (Fig 4) within the brain. As contrast for the micro-CT is provided by eosin and phase contrast processing, the processing for the presented protocol does not preclude further histological sectioning and imaging of the brain (Fig 2).

This work builds upon the prior protocol, developed for systems neuroscientific questions, that uses osmium as the contrast agent for micro-CT^8^, which allows for use with commercially available micro-CT machines. Our protocol offers three key advantages over that previous protocol for systems neuroscience. First, our protocol does not require multiple hazardous processing steps such as osmication, dehydration and resin embedding—as described in Masís et al.^8^—replacing it with a single step of 5% eosin in water. Second, it allows for brain imaging with a partially intact skull, permitting the direct measurement of skull landmarks relative to brain structures, and if necessary, subsequent micro-CT of the eosin-stained brain without a skull (data not shown). Third, our protocol greatly simplifies the subsequent processing for histology as the tissue can be cut and stained immediately after micro-CT with a range of standard histological stains. Our microscopic analysis confirmed well-preserved neural morphology after incubation in 5% eosin (Fig 2). Our protocol does, however, benefit from access to a synchrotron source to take full advantage of phase contrast processing. Although, there are a limited number of synchrotrons, they are available to all. Recent work has shown that phase contrast processing could be possible on commercially available micro-CTs^13^, allowing for future studies to potentially relax the need for a synchrotron source.

In developing the presented method, we sought to keep the protocol simple. Given the rapid advancement of micro-CT technology and processing, it is entirely possible that even further simplifications to the protocol could occur. Our goal was to use the method as a means towards neuroanatomic evaluation for systems neuroscience experiments. Given our ability to evaluate lesions and electrode placement, this method has achieved that goal. As such we have not compared our method to other plausible options, such as those using different staining techniques. Further studies can explore such comparisons.

Our protocol, while applied to the rat brain with a focus on the hippocampus, is compatible with other species and other brain regions. With the expansion of organisms used for genetic manipulations, neural recordings, and behavior this protocol can be very useful for all stages of the process. Our protocol also presents an excellent way to quickly generate a stereotactic atlas for guiding further experimentation. We note however, that micro-CT has limitations on the maximal field of view available, making this methodology less appropriate for larger brains.

There have been tremendous advances in recording technology to extract precise quantitative physiological information from many regions and cells throughout the brain^43-45^. However, the way in which we evaluate the precise locations at which those devices recorded often rests on descriptive and qualitative metrics. Given the ever-expanding corpus of precise anatomical information, there is a need to improve the precision and quantitation of electrode locations, and our protocol provides such an approach. It also allows for rapid and more objective quantification of lesion extent and other features that one might wish to compare across brains.

## Acknowledgements

We thank J Masis and A Joshi for helpful discussions. We thank EL Denovellis, H Barnard and A MacDowell for technical assistance. This work was supported by grants from the Jane Coffin Childs Memorial Fund for Medical Research (DBK), the UCSF Physician Scientist Scholars Program (DBK), an NIH R25 (R25MH060482) (DBK), an NIH Core Grant for Vision Research (NIH-NEI EY002162) (LDS), Research to Prevent Blindness (RPB Unrestricted Grant to the UCSF Department of Ophthalmology) (LDS) the Simons Foundation for Autism Research grant (291584) (LMF) and the Howard Hughes Medical Institute (LMF). This research used resources of the Advanced Light Source, a DOE Office of Science User Facility under contract no. DE-AC02-05CH11231.

## Author contribution

DBK, VK and DYP conceived of the study. DBK, VK and DYP developed the protocol. DBK and DYP collected the data. DBK, RN, ZY and CS performed the surgeries. DBK, RN, LDS and LMF developed the analyses. DBK, RN, DAM, ZY, and GMW analyzed the data. DBK, VK, DYP and LMF wrote the paper.

## Conflict of interest

VK is employed (part-time) by Neuralink Corp. The other authors declare no conflicts of interest.

**Video 1:** Video showing the entire three-dimensional rendering of the same brain from Fig 1 and virtually slicing the brain along the sagittal, coronal and horizontal axes.

**Supplementary Figure 1:**
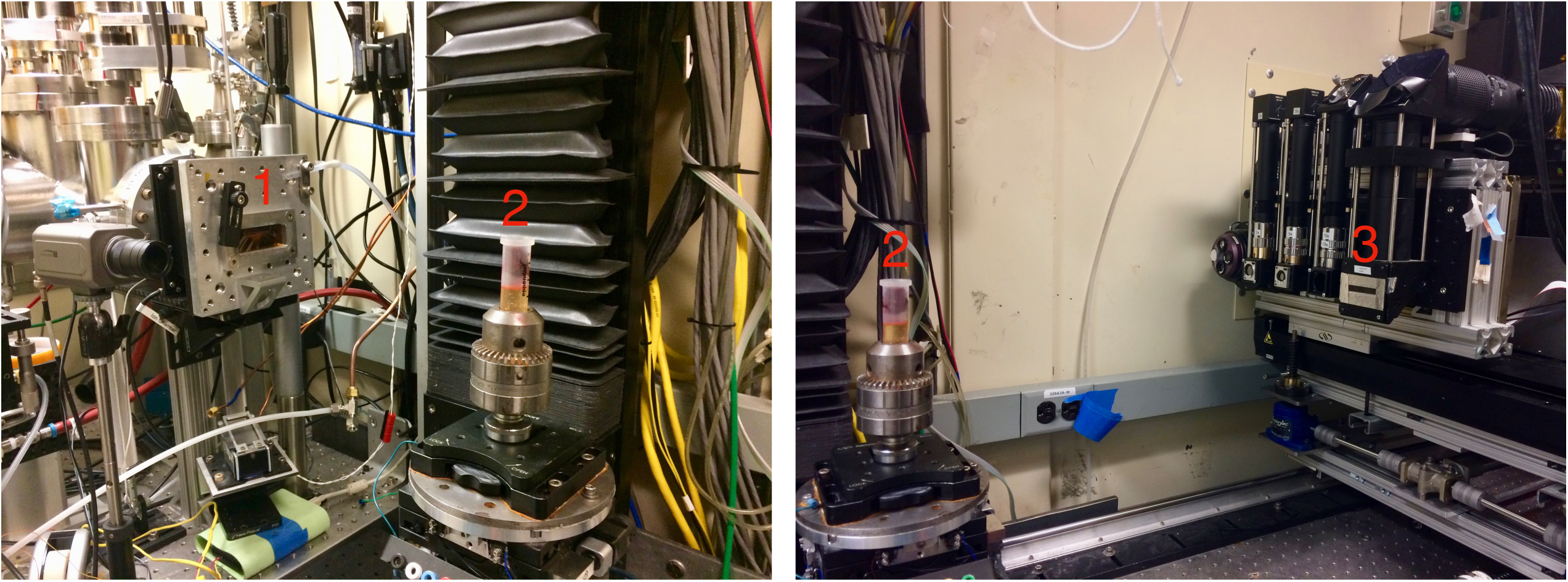
Layout of micro-CT at the ALS. 1 indicated the location of the x-ray source. 2 indicated the location of the sample, which rotates. 3 indication the location of the collector optics.

**Supplementary Figure 2:**
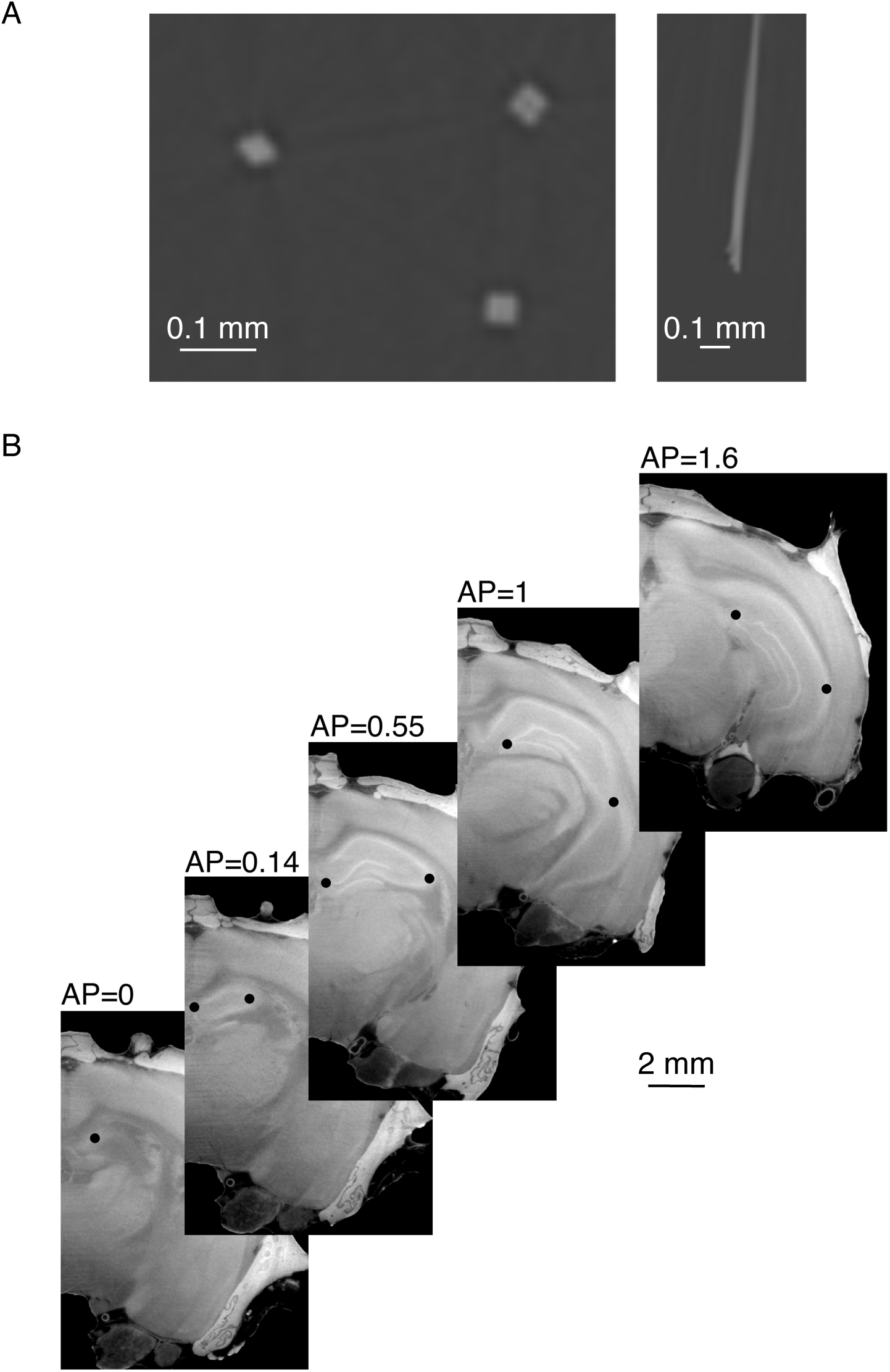
A) Fine scale tetrode structure can be visualized with the micro-CT. Left shows a virtual horizontal slice, where the four individual electrodes that make up a single tetrode can be seen. Right show a virtual coronal slice, where the individual electrode and the braiding of the tetrode can be seen. B) Example brain showing the location of the coordinates the make up the two-dimension space from Fig 4C. Virtual coronal slices of an example brain, with sample points of the two-dimensional space demarcated as black dots. At AP=0 the dot shows the origin point. At AP=1 the rightmost dot shows the point (1,1). For the rest of the slices the rightmost dot shows the medial part of the dentate cell layer and the leftmost dot shows the lateral aspect of the hippocampal cell layer. Scale bar applies to all images within the panel.

